# Inferring the Demographic History of Japanese Eel (*Anguilla japonica*) from Genomic Data: Insights for Conservation and Fisheries Management

**DOI:** 10.1101/2021.11.18.468312

**Authors:** Leanne Faulks, Prashant Kaushik, Shoji Taniguchi, Masashi Sekino, Reiichiro Nakamichi, Yuki Yamamoto, Hiroka Fujimori, Chiaki Okamoto, Sakie Kodama, Ayu Daryani, Angel Faye Manwong, Ishmerai Galang, Noritaka Mochioka, Kiyo Araki, Motoo Suzuki, Yoshitsugu Kaji, Takumi Ichiki, Tetsuya Matsunaga, Hiroshi Hakoyama

## Abstract

1. Assessing the status or population size of species is a key task for wildlife conservation and the sustainable management of harvested species. In particular, assessing historical changes in population size provides an evolutionary perspective on current population dynamics and can help distinguish between anthropogenic and natural causes for population decline.
2. Japanese eel (*Anguilla japonica*) is an endangered yet commercially important catadromous fish species. Here we assess the demographic history of Japanese eel using the pairwise and multiple sequentially Markovian coalescent methods.
3. The analyses indicate a reduction in effective population size (*N*_*e*_) from 38 000 to 10 000 individuals between 4 and 1 Ma, followed by an increase to 80 000 individuals, between 1 Ma and 22−30 kya. Approximately 22−30 kya there is evidence for a reduction in *Ne* to approximately 60 000 individuals. These events are likely due to changes in environmental conditions, such as sea level and oceanic currents, especially around the last glacial maximum (19−33 kya).
4. The results of this study suggest that Japanese eel has experienced at least two population bottlenecks, interspersed by a period of population growth. This pattern of demographic history may make Japanese eel sensitive to current and future population declines. Conservation management of Japanese eel should focus on practical ways to prevent further population decline and the loss of genetic diversity that is essential for the species to adapt to changing environmental conditions such as climate change.

## Introduction

Assessing the status or population size of species is a key task for wildlife conservation and the sustainable management of harvested species. However, one-time static estimates of population size provide a limited amount of information; assessing changes in population size over time yields greater insight into population dynamics. For example, understanding historical changes in population size places current population decline in an evolutionary context and can help distinguish between anthropogenic and natural causes for population decline (Dong et al., 2021). In addition, changes in population size have predictable effects on genetic variation. Bottlenecks, drastic reductions in population size over a short period of time, result in the loss of genetic variation and if the original population size was small or the population growth rate is low, recovery of genetic variation is slow (Nei et al., 1975). In such populations, sustained low levels of genetic diversity can reduce population fitness and limit the ability to adapt to environmental changes (Reed & Frankham 2003). Understanding this ‘genetic background’ of a species that is experiencing current population decline can indicate the susceptibility of that species to further genetic deterioration or even extinction (Abascal et al., 2016). Therefore, assessing changes in population size over historical timescales, termed demographic history, is applicable to contemporary conservation and management.

The demographic history of species can be inferred by estimating the effective population size (*N*_*e*_; the size of an ideal population that experiences genetic drift at the same rate as the observed population) of the species at various points in time. One approach is to use the pairwise and multiple sequentially Markovian coalescent (PSMC and MSMC) models to estimate *N*_*e*_ from whole-genome sequences derived from just a single individual (Li & Durbin 2011). This approach has even been successful for ancient DNA samples, including extinct species. Studies of the demographic history of extinct species observe two main patterns: fluctuations in population size, e.g., passenger pigeon (Hung et al., 2014), scimitar-horned oryx (Humble et al., 2020), and woolly rhinoceros (Lord et al., 2020); and gradual population reduction over time, e.g., Steller’s sea cow (Sharko et al., 2021) and two New Zealand passerines (Dussex et al., 2019). Similar patterns of demographic history have also been observed in some studies of threatened species, e.g. fluctuations: Iberian lynx (Abascal et al., 2016), Sumatran rhinoceros (Mays Jr et al., 2018), greater bamboo lemur (Hawkins et al., 2018), and red pandas (Hu et al., 2020); gradual reduction: brown hyena (Westbury et al., 2018); and long term low population: vaquita (Morin et al., 2021), African wild dog (Campana et al., 2016), and pygmy hog (Liu et al., 2021). Assessment of the demographic history of these threatened species is anticipated to contribute to their conservation management. However, these patterns of demographic history are not evident in all at risk species. For example, species that have experienced recent population declines, such as the commercially exploited fish species European eel (Nikolic et al., 2020), Atlantic cod and haddock (Tørresen et al., 2018), and yellowfin tuna (Barth et al., 2017), all show evidence of historical population size increase leading up until the last glacial maximum (LGM, 19−33 kya; Clark et al., 2009), after which they experienced population bottlenecks. Such bottleneck events may be considered as random catastrophes which can contribute to a species risk of extinction (Lande 1993). In these cases, it may be difficult for population size to recover given the combination of an LGM bottleneck and contemporary human-induced harvesting pressure. These examples highlight the importance of considering the demographic history of species in order to more fully understand current population dynamics and develop management strategies that are comprehensive and contribute to the long-term sustainability of species.

One commercially important species that has experienced recent declines in census population size attributed to fishing pressure, climate change, pollution, and habitat destruction (Tsukamoto et al., 2009), and is now listed as endangered by the IUCN, is Japanese eel (*Anguilla japonica*) (Pike et al., 2020; Tsukamoto et al., 2009). Japanese eel is a catadromous migratory species distributed across the rivers and estuaries of the northwestern Pacific, including Japan, China, Korea, Philippines, and Taiwan (Pike et al., 2020). After 5 to 15 years living in rivers and estuaries, yellow eels transform into silver eels and migrate to the spawning area near the West Mariana Ridge in the Pacific Ocean (Yokouchi et al., 2009). Following spawning the adults die and larvae drift on the North Equatorial and Kuroshio Currents where they metamorphose into glass eels and then arrive in waves along the east coast of Asia (Gong et al., 2019). Due to the complex life history and commercial importance of Japanese eel, there has been considerable research into its population genetic structure and population dynamics. Most studies on the population genetic structure of Japanese eel suggest that the species is panmictic (Gong et al., 2019, 2014; Han et al., 2010; Ishikawa et al., 2001; Minegishi et al., 2012; Sang et al., 1994). However, some studies have argued for the existence of low- and high-latitude groups (Chan et al., 1997; Tseng et al., 2006, 2009) and one recent study found evidence for the genetic differentiation of one population from the Kuma River, southern Japan (Igarashi et al., 2018). It is possible that these anomalous results are due to temporal fluctuations in genetic structure in an otherwise panmictic population, as also observed in European eel (Dannewitz et al., 2005; Pujolar et al., 2009). To date, studies of the demographic history of Japanese eel have found evidence for population size expansions 110−350 kya and 9−25 kya (Tseng et al., 2012), and a decline in population size 3.5−8 kya (Tseng et al., 2003). However, these inferences are limited in scope due to the use of few molecular markers (6−8 microsatellites, mtDNA D-loop sequences) and sampling from just a single location (1 river in Taiwan), so further analyses are warranted.

Determining *N*_*e*_ has been made easier and more accurate by the abundance of genomic data available, as well as the recent development of a variety of data analysis approaches (Fuentes-Pardo & Ruzzante, 2017; Spence et al., 2018). Investigating long-term historical changes in *N*_*e*_ can provide an evolutionary context within which to assess the nature of current population dynamics. This approach also improves our understanding of the response of species to past and future environmental change (Gattepaille et al., 2016). The overall aim of the current study was to assess the historical demography of Japanese eel. Inferences of past changes in *N*_*e*_ were made by utilising whole genome re-sequencing data of Japanese eel from two sources: the previously published data of Igarashi et al., 2018 and new data described here. The results were compared with those from previous studies on Japanese eel using ‘traditional’ markers (Tseng et al., 2012, 2003), as well as with similar analyses in other *Anguilla* species (Barth et al., 2020; Nikolic et al., 2020), to provide fisheries management agencies with a broader perspective of the current status of Japanese eel populations.

## Materials and Methods

### Inference of population demographic history

Past changes in *Ne* were estimated using the pairwise and multiple sequentially Markovian coalescent (PSMC and MSMC) methods (Li & Durbin, 2011; Schiffels & Durbin, 2014). These approaches examine an individual’s whole genome sequence and estimate the time to the most recent common ancestor of windows along the genome based on the density of heterozygous sites within those regions (Li & Durbin, 2011; Schiffels & Durbin, 2014).

The model is then used to estimate ancestral population sizes. For these analyses, a previously published dataset of 84 whole genome re-sequenced Japanese eels (Igarashi et al., 2018) was obtained from the National Centre for Biotechnology Information (NCBI) database under BioProject PRJDB5707. Duplicate reads were removed from the dataset and mean read depth assessed for each sample. As genome coverage is known to influence the accuracy of P/MSMC, due to the potential for false negative calling of heterozygotes in low coverage data (PSMC manual), only samples with a mean read depth greater than 18 (Nadachowska-Brzyska et al., 2015) were considered for the analyses. This resulted in just one glass eel from Sagami River, Kanagawa, Japan (sample ID: DRR091902). Thus, to obtain additional data with higher mean read depths, whole genome re-sequencing was performed on ten adult eel samples collected from Kuma River, Kumamoto Prefecture, Japan in 2018 (May−September). This data is available upon reasonable request to the authors.

The ten specimens from Kuma River were kept at −20°C until DNA extraction. Genomic DNA was extracted from a small piece of muscle tissue by using a Maxwell RSC Blood DNA Kit with a Maxwell RSC Instrument (Promega). A shotgun DNA library was constructed using a TruSeq Nano DNA High Throughput Library Prep Kit (Illumina), and the resulting library was subjected to pair-end sequencing with a Novaseq 6000 sequencer (Illumina) by Macrogen Japan. Trimmomatic v0.39 (Bolger et al., 2014) was used for adapter trimming and initial filtering (SLIDINGWINDOW, threshold phred-scale quality score of 25 within a window size of 4 bases; LEADING and TRAILING, 20; MINLEN, 50). The resulting pair-end reads were mapped onto a reference genome sequence of Japanese eel (Nakamura et al., 2017) using BWA v0.7.17 (Li 2013) with default parameters (the bwa-mem algorithm). Samtools v1.7 (Li et al., 2009) was used for additional filtering. The following were removed: reads with a mapping quality less than 10 (the subprogram, view); reads with an XA or SA tag (alternative hits and chimeric reads, respectively; standard grep command); unmapped reads (fixmate); and PCR duplicates (markdup).

Reads from the 11 samples were aligned to the draft reference assembly of Japanese eel (Nakamura et al., 2017) using BWA (Li 2013) and the consensus sequence for each was called using the mpileup command in bcftools (Danecek et al., 2021). Minimum base and mapping quality thresholds were set to 20. The total number of single nucleotide polymorphisms (SNPs) across the genome was estimated by vcftools (minQ 20, minDP 15, maxDP 120, max-missing 0.9) (Danecek et al., 2011). For the PSMC, variants were called using vcfutils.pl, setting the minimum and maximum read depths to one third and twice the mean depth of each sample, respectively. Then the PSMC package was used to convert files from vcf to psmc format and the analysis was performed with the following settings: iteration length 30, *T*_max_ 15, initial *θ*/*ρ* 5, and atomic time interval 64 (4+25*2+4+6). For the MSMC, the data was un-phased, so analyses were performed separately for each individual. Variant and input files were created for each individual using the MSMC tools scripts bamCaller.py and generatemulithetsep.py. The MSMC method requires separate input files for each chromosome, or in this case scaffold. Scaffolds that were longer than N75 (115 160 basepairs) were selected, resulting in 1719 input files per individual. The MSMC was then performed with default settings.

### Parameter Estimation and Plotting

To be scaled into time in years, the PSMC and MSMC output requires estimates of the species generation time and genome-wide mutation rate. A review of life history studies of Japanese eel indicated that silver eels (eels that are ready to undergo the spawning migration) range in age from 4 to 17 years, with a mean of 8 years for females (Han et al., 2009; Kotake et al 2007, 2005; Yokouchi et al., 2009). The spawning migration is estimated to take 6 months (Chang et al., 2016; Han et al., 2009), thus a generation time of 8.5 years was used to convert the PSMC and MSMC output to real time. The genome-wide mutation rate of Japanese eel was calculated by comparison with the European eel, *Anguilla anguilla*. The reference genome of European eel was obtained from the NCBI database under BioProject PRJNA561979. Each chromosome (1−19 and the mitochondrial genome) was aligned to the Japanese eel draft assembly using LASTZ (Harris, 2007). The number of matches and mismatches between the sequences of the two species were counted and then summed across all chromosomes. The mutation rate (per nucleotide per year) was then calculated as *μ* = (number of mismatches/total length)/2*t*, where *t* is the divergence time between Japanese and European eel. The divergence time was estimated to be 13.6 million years (Santini et al., 2013). Thus, *μ* = 8.7 × 10^−9^. The mutation rate per generation was then calculated as *μ* × 8.5, resulting in a mutation rate of 7.4 × 10^−8^ per nucleotide per generation. The results for the PSMC and MSMC analyses were scaled to real time using the values described above. Changes in *N*_*e*_ over time, for each individual and for the mean of all 11 individuals, were plotted by using R 4.1.0 (R Core Team, 2021).

## Results

Following the alignment of the 11 samples to the Japanese eel draft genome, 2 373 329 SNPs were identified. The 10 re-sequenced individuals had mean read depths of 42−59. The PSMC results indicated that the *Ne* of Japanese eel was stable at around 8 000 (7 700−8 400) individuals 1 Ma and then from approximately 500 kya *N*_*e*_ started to increase (Figure 1a, b). This population size increase continued until it peaked at approximately 88 000 (68−102 000) individuals around 30−40 kya. There was then a sudden reduction in *N*_*e*_ to approximately 63 000 (46−80 000) individuals around 30 kya. A population increase to approximately 74 000 (54−94 000) individuals occurred around 17 kya. The MSMC results indicated an initial reduction in *N*_*e*_ from 38 000 (32−60 000) to 10 000 (10−13 000) individuals between 1 and 4 Ma (Figure 1a, c). This was followed by a steady increase in *N*_*e*_ to approximately 81 000 (64−105 000) individuals around 22−32 kya. There was then a sudden reduction in *N*_*e*_ to approximately 56 000 (45−77 000) individuals around 22 kya. A population increase to 97 000 (60−178 000) individuals occurred around 14 kya.

**Figure 1.**
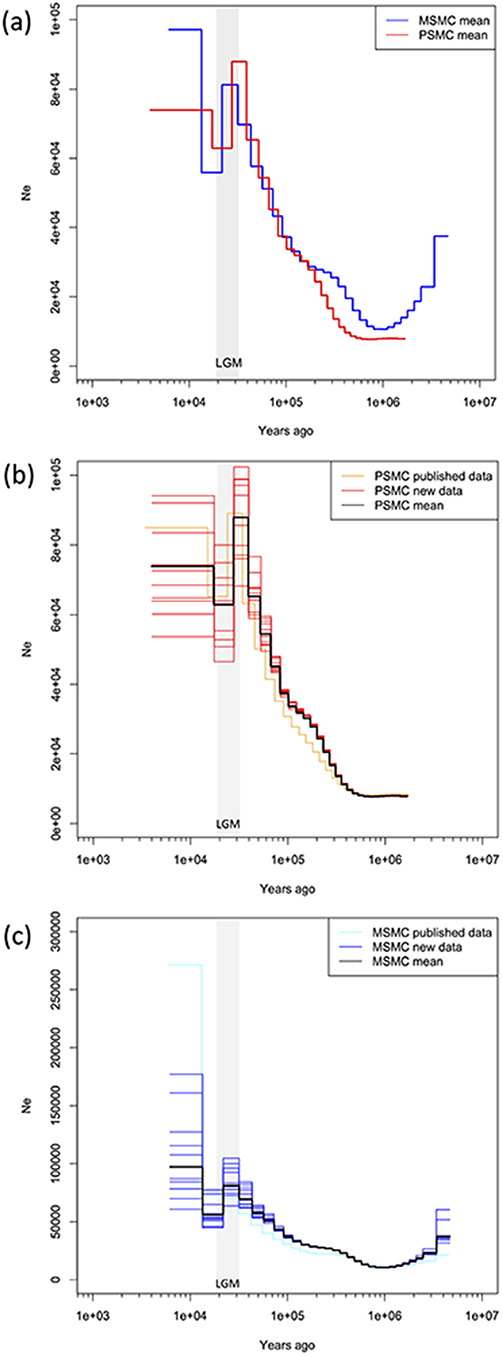
Plot of the changes in effective population size *N*_*e*_ over the last 10 My for Japanese eel. Effective population size was estimated from the whole genome re-sequencing data of 11 individuals by using the pairwise and multiple sequentially Markovian coalescent (PSMC and MSMC) methods. (a) Plot presenting the mean values of *N*_*e*_ for all 11 individuals analysed, PSMC = red, MSMC = blue; (b) Plot representing the PSMC results for 10 individuals sequenced in this study (red lines), 1 individual from previously published data (orange line), and the mean (black line); (c) Plot representing the MSMC results for 10 individuals sequenced in this study (blue lines), 1 individual from previously published data (light blue line), and the mean (black line).

The PSMC and MSMC inferences were very similar, although MSMC was able to infer *Ne* further back in time (5 Ma vs. 2 Ma). In addition, the inferences based on one individual from previously published data (Figure 1 b, c, yellow lines) were similar to those based on the data from 10 re-sequenced individuals obtained in this study (Figure 1 b and c, red and blue lines, respectively).

## Discussion

### Demographic history of Japanese eel

This study provided valuable insights into the demographic history and *N*_*e*_ of Japanese eel, a species of conservation and fisheries management concern. In summary, the results of this study provide the following general picture of the demographic history of Japanese eel: population reduction between 1 and 4 Ma, population growth between 30 and 800 kya, sudden population reduction around 25 kya (during the LGM), followed by population growth around 15 kya (post-LGM). Interestingly, the findings of this study were similar to those of a previous study that used mtDNA sequences (Tseng et al., 2012), suggesting that ‘traditional’ methods remain relevant and informative, especially in cases where the availability of funds, equipment, expertise, or high-quality DNA may prevent the use of NGS techniques. In addition, our analyses demonstrate the potential for publicly available genome data to contribute to conservation and fisheries management. The long-term view of population dynamics presented here helps provide some perspective on current population declines.

The MSMC inferred a reduction in the *N*_*e*_ of Japanese eel between 1 and 4 Ma, during the Pliocene. During this time, the formation of the Isthmus of Panama, together with the closure of the Indonesian seaway, caused a general shift in global oceanic circulation and a drop in sea surface temperatures that led to glaciation in the northern hemisphere (Cane & Molnar, 2001; Gallagher et al., 2015). It is also postulated that at this time the Kuroshio current did not reach the Japan/East China Seas, and that the Tsushima current inflow was weak and periodical until about 1.7 Ma (Gallagher et al., 2015). These factors may have restricted the larval drift and/or migration of Japanese eels, leading to declines in *N*_*e*_.

Both the PSMC and MSMC inferences indicated that the *N*_*e*_ of Japanese eel started to increase around 0.5−1 Ma. Notable changes to the oceanic environment in the western Pacific occurred during this period. Firstly, global sea levels were relatively high and large areas of south-eastern China and the Ryukyu Islands (southern Japan) were inundated, influencing the circulation of ocean currents (Kimura, 2000). Secondly, stronger glacial/inter-glacial cycles intensified the Kuroshio Current and the North Pacific Gyre (Gallagher et al., 2015). Together these conditions may have facilitated a population size expansion in Japanese eel, which continued until approximately 30 kya. The reduction in *Ne* that occurred around 30−22 kya is most likely due to changes in sea level, ocean current patterns, and sea surface temperatures associated with the LGM (19−33 kya). For example, during the LGM a land bridge between Taiwan and the Ryukyu Islands deflected the Kuroshio Current eastwards (Ujiié & Ujiié, 1999), thereby preventing the drift of Japanese eel larvae northwards to the rivers and estuaries of the Japanese coastline. Because the PSMC and MSMC methods perform less well for inferences of demographic history within the last 20 ky (Mather et al., 2020), we interpret population size changes after the LGM with caution. However, the results of this study (population growth 14−17 kya), a previous study of Japanese eel (population expansion 9−25 kya; Tseng et al., 2012), and studies of other species (three tropical *Anguilla* species; Barth et al., 2020), as well as other northwest Pacific fishes (Lu et al., 2020; Xu et al., 2019), all provide evidence for post-LGM population growth, suggesting that oceanic conditions were generally suitable for population expansion at that time. In the case of Japanese eel, perhaps abrupt sea level rise at the end of the LGM (14.5 kya; Clark et al., 2009) and the inundation of the Taiwan−Ryukyu land bridge 10 kya (Ujiié & Ujiié, 1999), which restored the Kuroshio Current to its present-day path, provided an opportunity for geographic population expansion and population size increase. We recommend further studies to clarify changes in *N*_*e*_ of Japanese eel within the last 20 ky, and to estimate the current *N*_*e*_.

### Implications for conservation and fisheries management

The results of this study suggest that, on an evolutionary time scale, Japanese eel may have experienced two population bottlenecks interspersed by a period of population growth. One previous study also detected evidence of a population bottleneck 3−8 kya (Tseng et al., 2003), a time period not covered by the PSMC or MSMC analyses. Finally, comparison to analyses of population demographic history in other *Anguilla* species suggest that Japanese eel has always had a relatively lower *N*_*e*_ compared to its congeners (Barth et al., 2020; Nikolic et al., 2020). Together, these findings suggest that Japanese eel has a demographic history of population size fluctuations and has experienced population bottleneck events. The genetic consequences of this demographic history may include reduced levels of genetic diversity and loss of adaptive potential that may make Japanese eel sensitive to current and future population declines. The results of this study demonstrate how climate induced changes in the pattern and intensity of ocean currents, particularly the Kuroshio and Tsushima currents, and consequent shifts in the availability of suitable habitat, led to changes in the historical *N*_*e*_ of Japanese eel. It is likely that these factors will also be important for the natural and long-term maintenance of *N*_*e*_ in the future. However, we acknowledge that it is difficult to predict the influence future climate change will have on Japanese eel population size because of the interaction between the effects of changes in ocean currents and sea surface temperatures, and the uncertainty surrounding the degree to which other anthropogenic activities like habitat degradation and harvesting will continue to contribute to population decline.

In conclusion, this study found that Japanese eel has experienced at least two population bottlenecks throughout its evolutionary history, a genetic background that warrants caution. Conservation and fisheries management of Japanese eel should focus on practical ways to prevent further population decline and the loss of genetic diversity that is essential for the species to adapt to changing environmental conditions such as climate change. Finally, we wish to highlight the valuable perspective that can be gained by assessing historical population demography and hope that this and similar studies continue to contribute to conservation and fisheries management.

## Acknowledgements

This work was supported by the Research and assessment program for fisheries resources, the Fisheries Agency of Japan. We thank Dr. Mayumi Kobayashi for helpful comments. We have no conflict of interest to declare.

## Notes

### Competing Interest Statement

The authors have declared no competing interest.

